# Laboratory horror stories: Poison in the agars

**DOI:** 10.1101/2024.06.06.597796

**Authors:** Mari K. Davidson, Reine U. Protacio, Dominique Helmlinger, Wayne P. Wahls

## Abstract

The fission yeast *Schizosaccharomyces pombe* is a single-celled eukaryote that can be cultured as a haploid or as a diploid. Scientists employ mating, meiosis, and the plating of ascospores and cells to generate strains with novel genotypes and to discover biological processes. Our two laboratories encountered independently sudden-onset, major impediments to such research. Spore suspensions and vegetative cells no longer plated effectively on minimal media. By systematically analyzing multiple different media components from multiple different suppliers, we identified the source of the problem. Specific lots of agar, from different suppliers, were toxic. Interestingly, the inhibitory effect was attenuated on rich media. Consequently, quality control checks that use only rich media can provide false assurances on the quality of the agar. Lastly, we describe likely sources of the toxicity and we provide specific guidance for quality control measures that should be applied by all vendors as preconditions for their sale of agar.

**Graphical Abstract:** 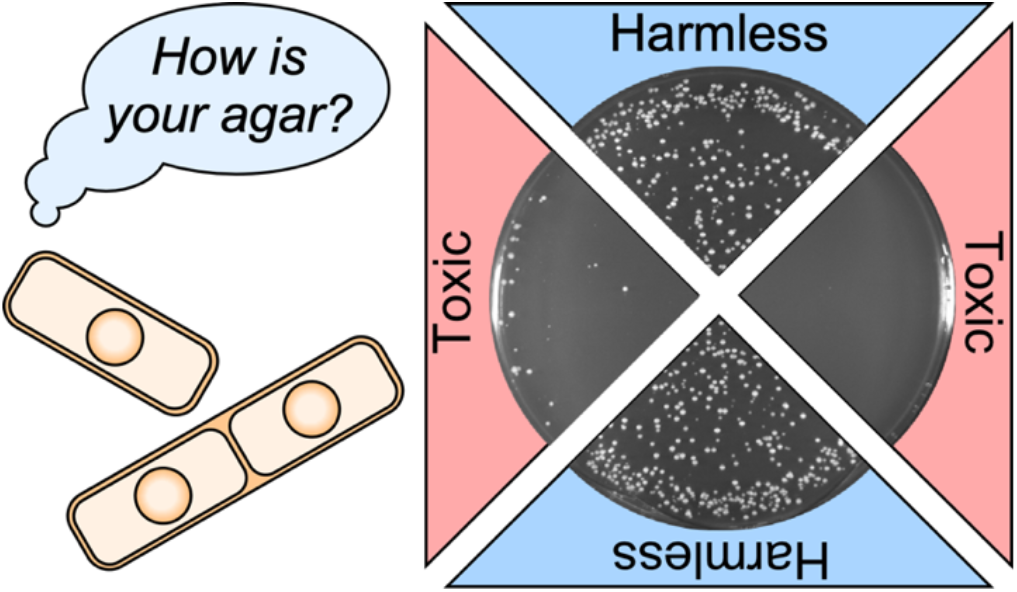

**Take-away:**

- Sporadically, batches of agar from different suppliers strongly inhibit the plating efficiency of *S. pombe* spores and vegetative cells on minimal media.
- Quality control checks that are not quantitative or that use only rich media can provide false assurances on the quality of the agar.
- Vendors should conduct rigorous, thorough, organism-specific tests for potential toxicity of each lot of agar as a pre-condition for its sale.

## Introduction

Agar is a hydrocolloidial gelling agent that is extracted from the cell walls of red algae (phylum *Rhodopyta*), often from numerous different species in the genera *Gracilaria* and *Gelidium* (which together have more than 500 recognized species) (Borg et al., 2023; Porse & Rudolph, 2017; Vandepitte et al., 2018). Agar is prepared by a crude extraction process that includes hot water extraction, gelling, drying and milling; refinements often include alkaline hydrolysis to improve yield and quality. Agar is composed primarily of agarose (heterogeneous polymers of the disaccharide agarobiose) and agaropectin (a heterogeneous mixture of galactan polymers). It is a particularly useful gelling agent because it displays hysteresis, with a gelling temperature around 40 °C and a melting temperature around 85 °C.

The primary use of agar is in the food industry, where it is employed to stabilize, thicken, gel, or otherwise modify the textural properties of food items; it also provides dietary fiber, making it a useful digestive aid (Bose et al., 2023). The second most common use for agar, which consumes about 10% of the world’s supply (Kim et al., 2017; Porse & Rudolph, 2017), is to solidify culture media for the isolation, cultivation and analyses of microorganisms (Koch, 1882). Additional biological and biomedical applications include cell motility assays (Partridge & Harshey, 2013); plant propagation (Ma et al., 2023), drug screening (Bidaud et al., 2021); and analyzing the invasive potential of cancer cells (Wang, 2023). Agar can also be used in hydrogel films (Ma et al., 2023); for applications such as wound dressings (Shen et al., 2021); for environmental remediation like adsorption of heavy metals from waste water (Rani et al., 2018); as nanoparticles for drug delivery (Gaikwad et al., 2024); and as scaffolds for immunoassays (Hornbeck, 2017).

Hundreds of research laboratories use agar plates to propagate and study an outstanding model eukaryotic organism, the fission yeast *Schizosaccharomyces pombe* (Harris et al., 2022). This single-celled eukaryote is about as distant evolutionarily from the budding yeast *Saccharomyces cerevisiae* as either yeast is from humans (Heckman et al., 2001; Sipiczki, 2000), and many properties of fission yeast biology more closely resemble those of humans (Vyas et al., 2021). Fission yeast can be cultured as a haploid or as a diploid and these two states can be easily interconverted by mating and meiosis (Kawamukai, 2024; Ohtsuka et al., 2022). This supports powerful genetic approaches that are complemented by equally powerful molecular tools [e.g., (Billmyre et al., 2022; Gao et al., 2014; Ishikawa & Saitoh, 2023; Storey et al., 2019; Torres-Garcia et al., 2020)], which has made fission yeast an eminent model to study a variety of broadly conserved eukaryotic processes [see (Harris et al., 2022; Hoffman et al., 2015; Nurse, 2020; Rutherford et al., 2024) and refs therein]. For example, we use this model organism to discover mechanisms by which specific DNA sequences, their binding proteins, and chromatin remodeling factors control the positioning of meiotic recombination throughout the genome (Mukiza et al., 2019; Protacio et al., 2022; Storey et al., 2018). In our line of research—as in many other areas of research using diverse organisms—we rely heavily on being able to measure with precision the titers of viable cells.

Here, we report sudden-onset reductions in the plating efficiency of fission yeast spores and cells which had catastrophic impacts on the progress of our research programs. We describe how we traced that problem to sporadic toxicity within specific batches of agar from multiple different suppliers. We discuss likely sources for this toxicity and we provide specific guidance for quality control measures that should be applied by vendors as preconditions for their sale of agar.

## Materials and Methods

### Fission yeast strains and genotypes

The names and genotypes of *S. pombe* strains used in this study are: WSP 3776 (*h*^*-*^); WSP 5819 (*h*^*+*^ *ade6-M375*); WSP 7850 (*h*^*-*^ *ade6-3049*); and DHP 148 (*h*^*90*^). Spore suspensions were from crosses between WSP 5819 and WSP 7850.

### Culture media and methods

Standard formulations were used for synthetic minimal media (Edinburg minimal media, EMM), minimal media (nitrogen base, NB), rich media (yeast extract, YE), and sporulation media (SP) (Forsburg & Rhind, 2006; Gutz et al., 1974; Kon et al., 1998; Wahls et al., 1993). An “L” or “A” is included in the name to designate liquid or agar (solid) media, respectively (e.g., NBA is nitrogen base agar). When required to support the growth of adenine-auxotrophic strains, adenine was included in the minimal media at 100 µg/ml (Gutz, 1971). Standard methods were used for yeast culture, for genetic crosses to induce mating and meiosis, and for the preparation of ascospore suspensions (Gao et al., 2008; Kon et al., 1998). Numbers of cells or spores in suspensions were determined using a hemocytometer.

## Results

In this study we used three types of culture media that are broadly employed by the fission yeast research community. Our rich media was yeast extract liquid (YEL) or agar (YEA); for minimal media we used nitrogen base liquid (NBL) or agar (NBA), as well as Edinburgh minimal media liquid (EMML) or agar (EMMA). The only difference between the liquid and solid media of each type was the inclusion of 2% agar in the latter.

### Sudden-onset reduction in the plating efficiency of spores

In July of 2023, we encountered a major, puzzling impediment to our research. Plating a known number of spores on NBA media yielded extremely low colony titers (**Figure 1A**). This low efficiency of plating (EOP) suggested that a substantial fraction of the spores were inviable or that there was a problem with the media. Since we often keep stocks of spore suspensions from previous experiments, we had available stocks with known, high titers. When we plated those viable spores on the newer NBA media they plated inefficiently, suggesting that the problem was with the media, rather than the spores. Notably, the compromised plating efficiency was observed for multiple different genotypes of spores and for vegetative cells (described subsequently), further supporting the idea that there was a problem with the culture media.

**Figure 1.**
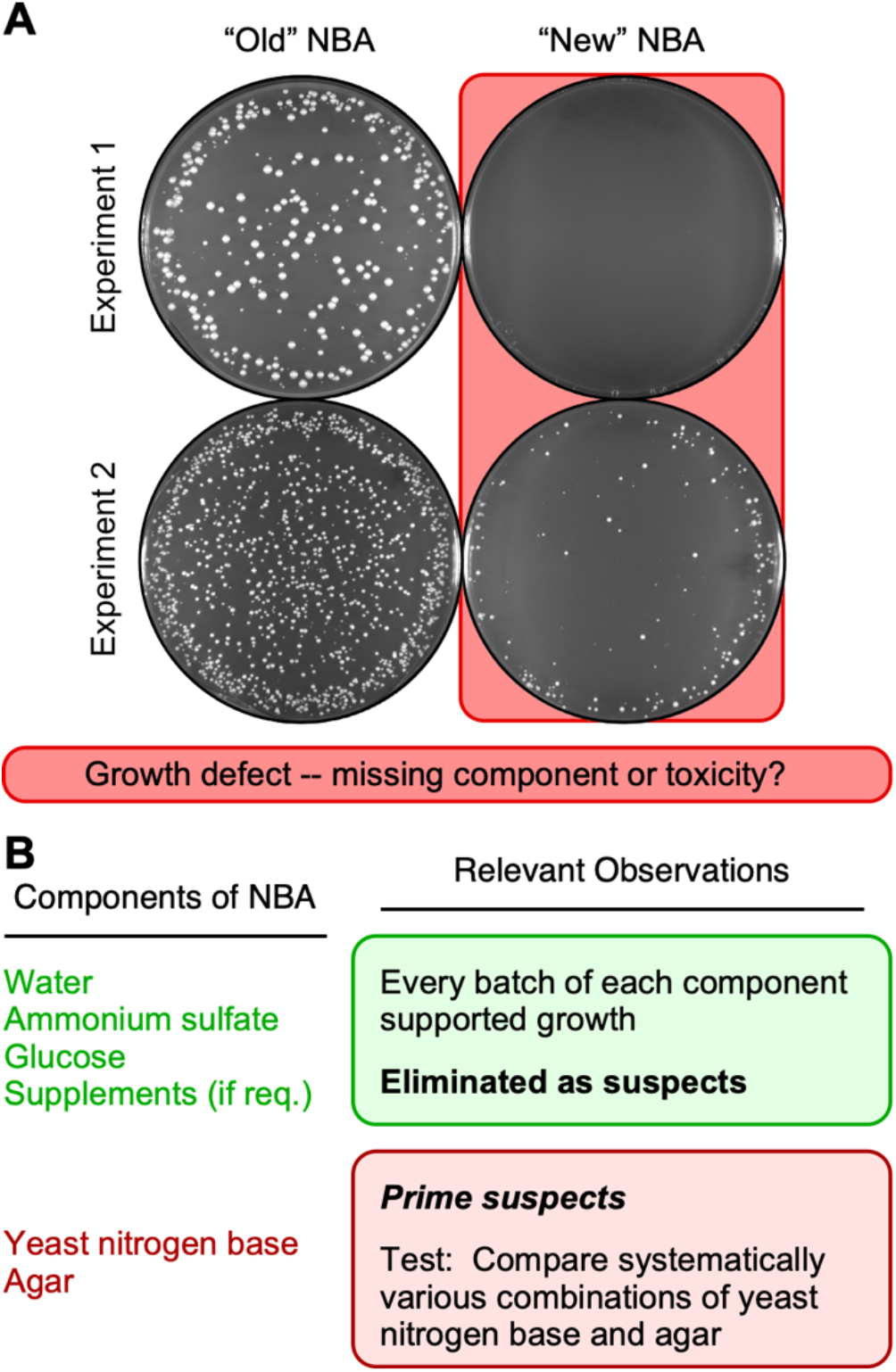
Spores plate less efficiently on some batches of NBA media than on others. **(A)** Images show representative results obtained with four different batches (i.e., separate preparations) of nitrogen base agar that were prepared contemporaneously. The batch labels (*old, new*) reflect the relative order in which the media components were purchased. Within each experiment, aliquots of the identical spore suspension were plated in parallel on two different batches of media (1,290 spores per plate in experiment 1 and 2,500 per plate in experiment 2). Background highlights (*coral* color) emphasize the lower EOP on one batch of media versus the other. **(B)** Rationale and approach. Diagram lists components of NBA media; empirical analyses of individual components focused attention on the ones most likely to be responsible for the media batch-specific differences in EOP.

To test whether there was a problem with the culture media, we plated spore suspensions in parallel on different batches of NBA that had identical formulations but contained media components purchased at different times or from different suppliers. The results were striking. Spores plated efficiently on our “old” NBA media, which contained components that we purchased in the past, whereas spores plated poorly on our “new” NBA media, which was made using more recently purchased components (**Figure 1A**). Similar results were obtained whether the “old” media had been stored for months as plates in the refrigerator or had been made freshly in parallel with the “new” media. Since the major differences in EOP occurred even when the biological samples were identical (e.g., **Figure 1A**), we conclude that the differences are attributable to the media itself. Presumably, some ingredient or component in the NBA media was missing or toxic.

### Narrowing down the list of suspects

We prepare our NBA minimal media (and other types of culture media) from its individual components (**Figure 1B**). We used two approaches to test whether a given component of the media was insufficient or toxic. First, if a specific batch of a given component was still available (e.g., bottle of ammonium sulfate solution), we retested that specific batch in combination with different batches of other components. Second, for a given component, we tested whether different lots or batches of that component (e.g., different bottles of ammonium sulfate) supported growth when used to make media in which the other components were maintained constant. These experiments revealed no problems with the water, ammonium sulfate, glucose and supplements used to make the media. Every lot and every batch of each component supported the efficient plating of spores and vegetative cells on solid media and the growth of cells in liquid cultures. We then applied a similar approach to test the prime suspects, yeast nitrogen base and agar.

### Low efficiency of spore plating is caused by the agar, not the nutrients

Since agar is a relatively inert gelling component of NBA media and the nutrients are provided by the nitrogen base (**Figure 1B**), it seemed likely that the differences in EOP were due to the batch or lot of nitrogen base. To test this hypothesis, we conducted a series of experiments in which we combined systematically different lots of nitrogen base and agar, changing only one variable at a time. Moreover, because we had historically purchased much of our nitrogen base and agar from one supplier, we compared in parallel multiple different lots from that vendor, as well as lots purchased from different vendors. Interestingly, each lot of nitrogen base from each supplier supported highly efficient plating of spores (e.g., **Figure 2**, left column). We conclude that each lot of nitrogen base contains all of the nutrients (other than carbon source) that are necessary and sufficient to sustain efficient spore germination and colony growth. We also conclude that those batches of nitrogen base contain no detectable inhibitors of growth. We therefore reject our hypothesis that the low EOP was caused by a defect or insufficiency in the nitrogen base (nutrients).

**Figure 2.**
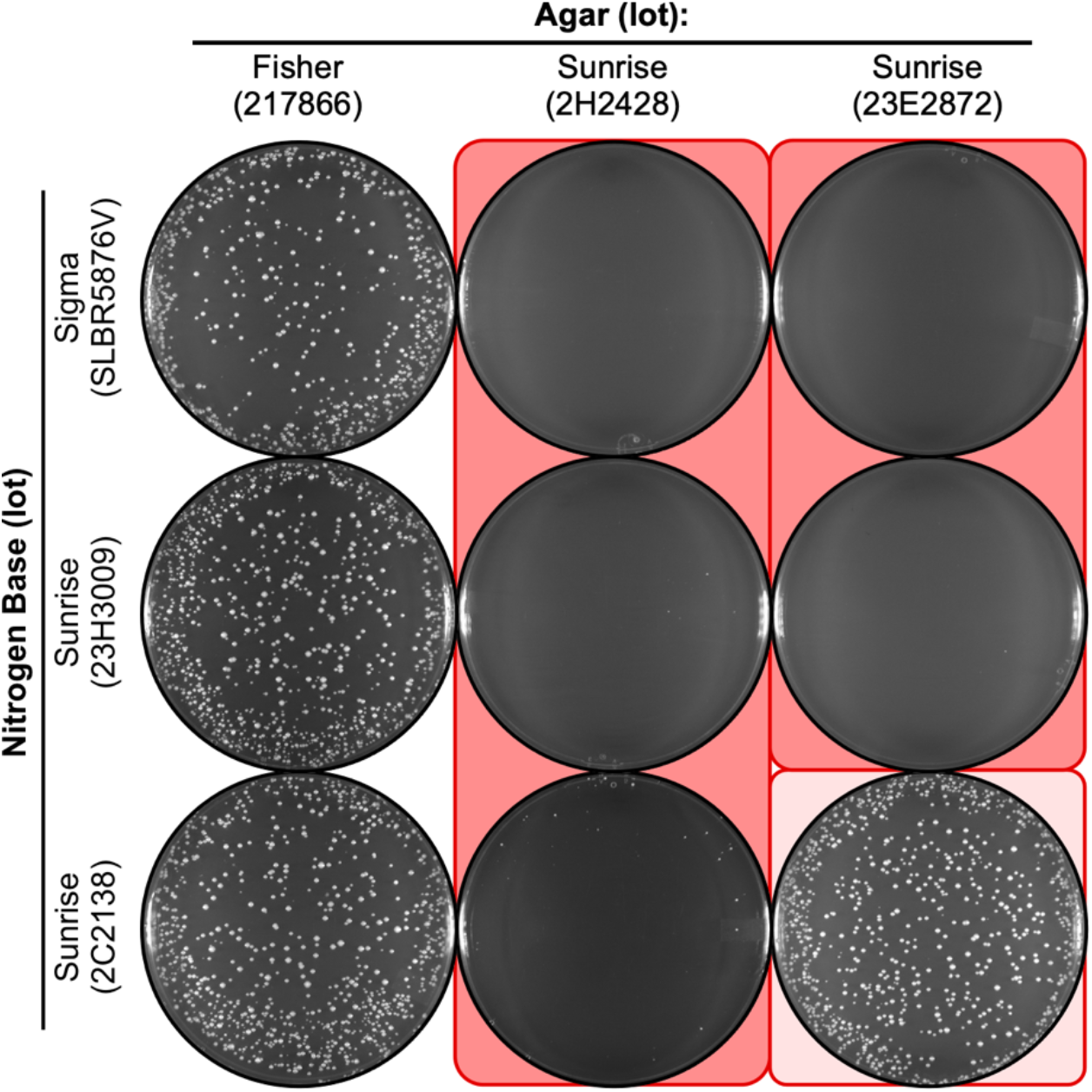
Agar lot-specific inhibition of spore colony growth on NBA. Nine different batches of NBA media were prepared using the indicated lots of nitrogen base and agar as variables. Images show EOP for representative platings of the same spore suspension (2,500 spores per plate) on the different media. Note that each lot of nitrogen base can support growth (column 1) and that a subset of the lots of agar (columns 2, 3) reduce the EOP. In this and subsequent figures, the degree of shading (*coral* color) reflects the magnitude of the inhibitory effect.

Notably, when the various lots of nitrogen base that were known to support growth were brought together with some lots of agar, the EOP fell dramatically (**Figure 2**, middle and right columns). Interestingly, other lots of agar (e.g., **Figure 2**, left column), including prior lots that we had purchased from our historically preferred vendor (e.g., **Figure 1A**, left column), did not inhibit the EOP. We conclude that there is an agar lot-specific inhibition in the EOP of fission yeast spores on the NBA minimal media. Since agar is a solidifying agent that provides no nutrients to the culture media, we can conclude that the inhibition is due to the presence of toxic agent(s) within the affected lots. Furthermore, because this toxicity was manifest for some but not all lots of agar from the same vendor (**Figure 1** and **Figure 2**), we conclude that it is a sporadic problem.

### Nutrients in the media can affect the toxicity of the agar

The experiments in which we systematically changed one variable at a time also revealed that the agar lot-specific toxicity displays variable penetrance. For example, agar from lot 23E2872 strongly suppressed the EOP of spores on NBA that contained nitrogen base from lots SLBR5876V and 23H3009 (**Figure 2**, right column). However, when the same agar was in combination with nitrogen base from lot 2C2138, spores plated efficiently (**Figure 2**, right column). A similar result was observed for agar from lot 2H2428: When in combination with nitrogen base from lots SLBR5876V and 23H3009, this agar strongly suppress the EOP; whereas the EOP was improved slightly when this agar was in media that contained nitrogen base from lot 2C2138 (**Figure 2**, middle column). Thus, some batches of nitrogen base can suppress the toxicity of the agar. Moreover, the same batch of nitrogen base can suppress the toxicity of multiple, different batches of agar. Interestingly, the degree of suppression varied in an agar lot-specific fashion (**Figure 2**, compare bottom plates, middle and right columns). This provides compelling evidence that some lots of agar are more toxic than other lots, which is concordant with our conclusion that some lots of agar are toxic and others are not. A similar logic applies for the suppression of toxicity by only some batches of nitrogen base (**Figure 2**, row 2 versus row 3), which is likely mediated by lot-to-lot differences in the abundance of the suppressive component(s) within the nitrogen base. The richness of the nutrients in the lot of nitrogen base is a good candidate for suppression of the toxicity (evidence described below).

We can summarize the preceding results and conclusions using a “poison and antidote” analogy. The effective amount of toxicity (“poison”) in the agar can vary from lot-to-lot; similarly, the amount of counteracting factor (“antidote”) in the nitrogen base can vary. Correspondingly, the EOP in the presence of the poison will be influenced by the relative amounts of both the poison and the antidote.

### A delicate balance between low EOP and high EOP

Several observations provide further insight into the nature of the variable penetrance of toxicity. First, for toxic agar lot 23E2872, different lots of nitrogen base from the same vendor differentially suppressed the toxicity (**Figure 2**, right column): Nitrogen base lot 23H3009 had a plating efficiency near zero, whereas lot 2C2138 supported nearly wild-type EOP. The fine balance can also be seen between lots of toxic agar (**Figure 2**, bottom row). When nitrogen base lot 2C2138 was together with agar lot 2H2428, the EOP was very low; when the same nitrogen base was in combination with agar lot 23E2872, the EOP was much higher. We conclude that the variable penetrance (low versus high EOP) is dictated by the proportionate amounts of the inferred toxin (agar lot) and the inferred counteracting agent (nitrogen base lot).

The agar lot-specific reduction in the plating efficiency of spores was reproducible between experiments where the media formulations and individual components were identical, but were prepared in separate batches (compare **Figure 3** to **Figure 2**). In each case there was also variable penetrance. However, there were informative differences in EOP between experiments for some combinations of agar lot and nitrogen base lot. For example, when toxic agars from lots 2H2428 and 23E2872 were with nitrogen base lot SLBR5876V, the EOP was either near zero (**Figure 2**) or about 10% (**Figure 3**). Similarly, when the toxic agar lot 2H2428 was with nitrogen base lot 2C2138, the EOP was near zero (**Figure 2)** or about 50% (**Figure 3**).

**Figure 3.**
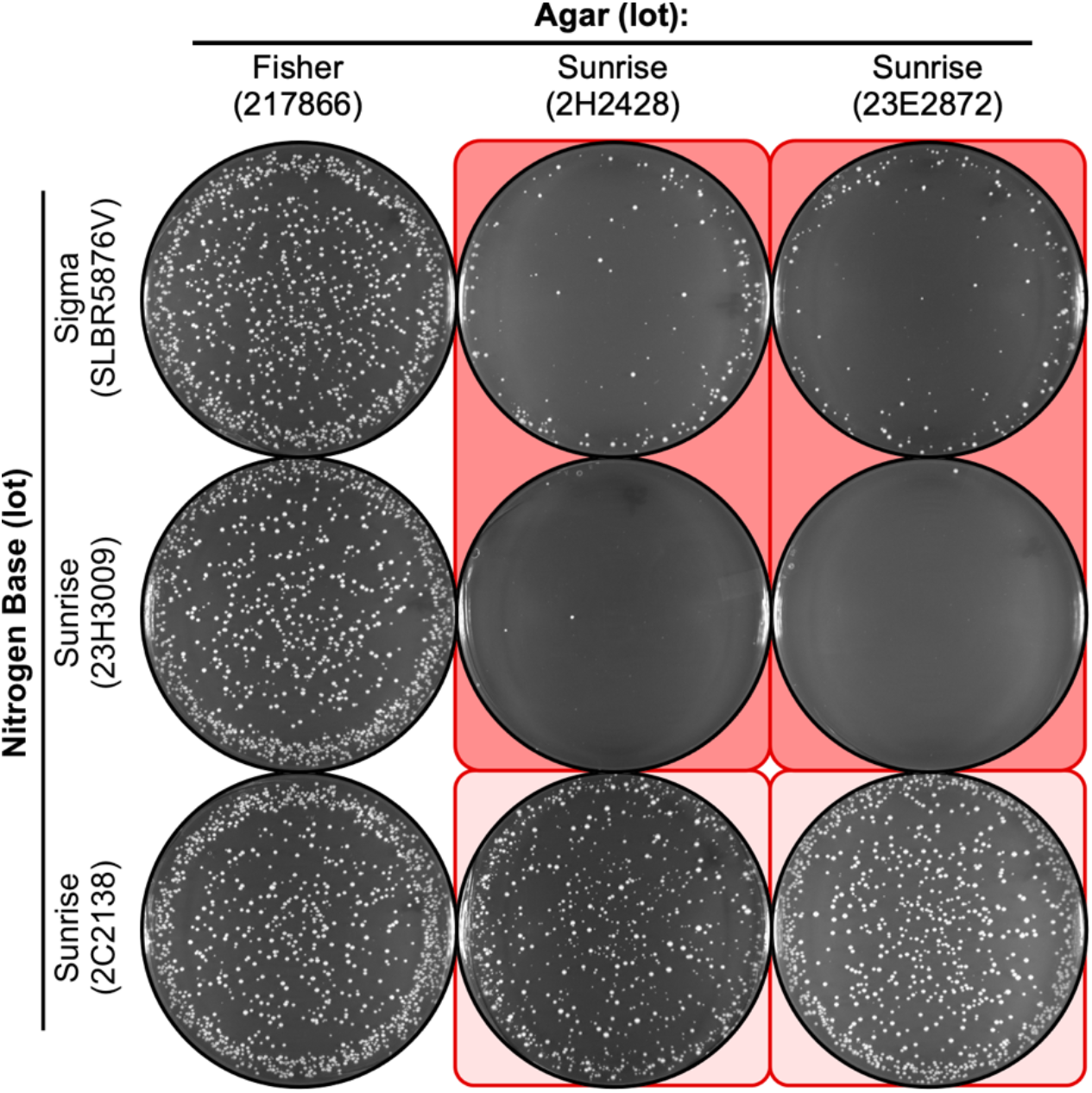
Reproducibility and variable penetrance batch-to-batch on NBA. The approach and media formulations were identical to those in Figure 2 but employed separately prepared batches of each media. Note that the same lots of agar reproducibly reduce the EOP (columns 2, 3). Also note that the magnitude of the inhibitory effect can vary from experiment to experiment. For example, the reduction in EOP for row 1, column 2 is less severe than the reduction in EOP for the same configuration in Figure 2.

We conclude that there is a delicate balance between the amounts of the inferred toxic agent in the agar and the inferred counteracting agent in the nitrogen base; moreover, there seems to be a fairly narrow range within which there is a change between low and high EOPs. This is particularly insidious because even minor perturbations—such as differences that might occur during the sterilization of two identically formulated batches of the same media—can demonstrably affect the plating efficiency (compare results in **Figure 2** versus **Figure 3**). And as illustrated nicely by results described thus far, this type of variability can confound efforts to diagnose the source of problems with culture media.

### Toxicity and variable penetrance also affect the plating of vegetative cells

The low EOP of spores on NBA media that contains toxic agar might be due to defects in spore germination or in subsequent growth of the vegetative cells. To distinguish between these two modes of action, we plated aliquots of cells from log-phase liquid cultures on the identical media that were used to measure the EOP of spores. The results were essentially identical: Each lot of nitrogen base from each supplier supported highly efficient plating of vegetative cells (**Figure 4**, left column); one lot of agar supported high EOP (**Figure 4**, left column); two lots of agar were toxic (**Figure 4**, right two columns); and some lots of nitrogen base differentially suppressed the toxicity of the agar (**Figure 4**, light and dark background shading). We conclude that the agar lot-specific toxicity affects vegetative cell growth, not just the plating efficiency of spores.

**Figure 4.**
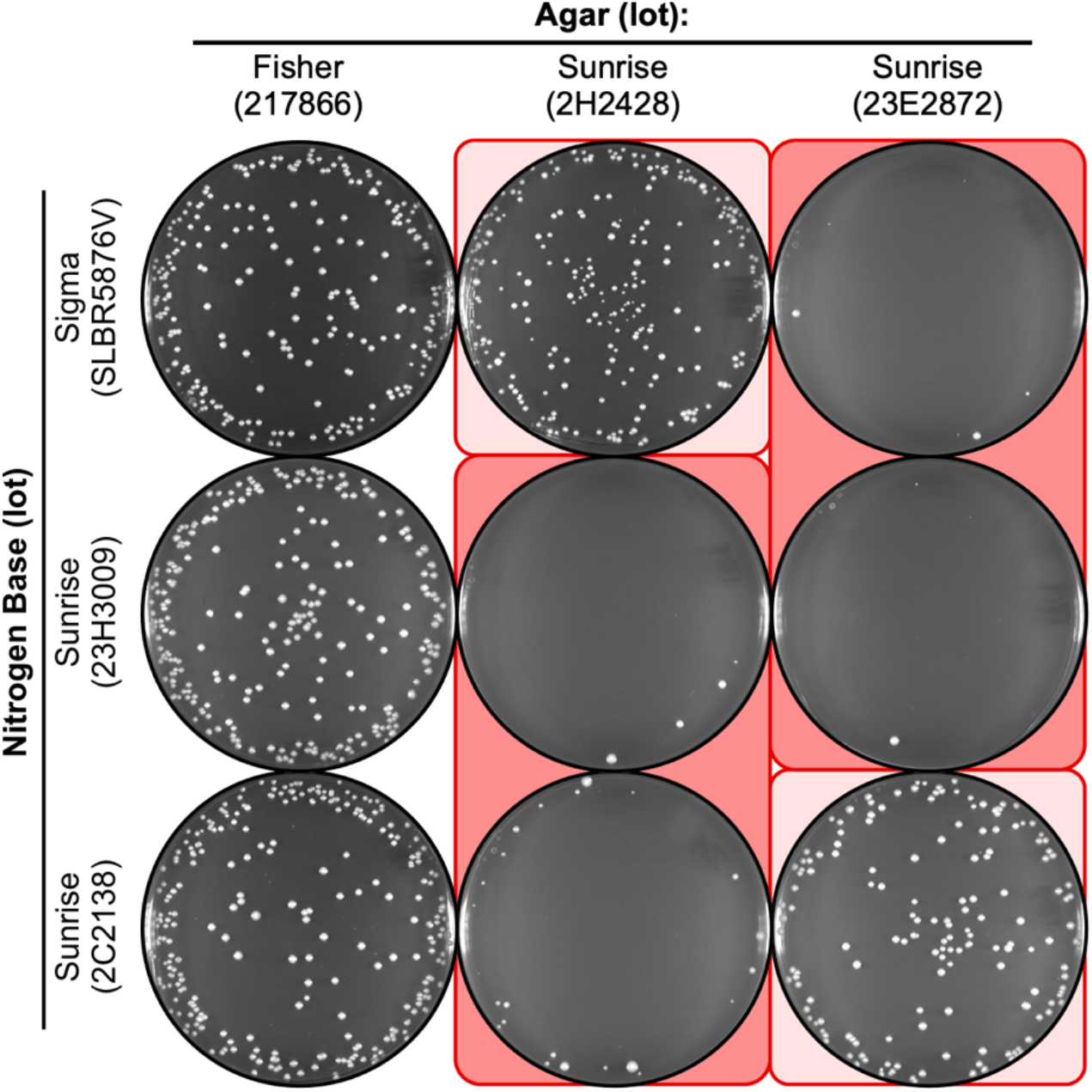
Agar lot-specific inhibition in plating of vegetative cells on NBA. Cells from log-phase growth in liquid culture were plated (using 400 cells per plate) on the indicated media. Note that each lot of nitrogen base can support growth (column 1) and that a subset of the lots of agar (columns 2, 3) reduce the EOP. The patterns of inhibition for vegetative cells are like those observed for the plating of spores (Figures 2, 3).

Similarly, we conclude that the principles and mechanisms for variable penetrance (proportionate ratios of “toxin” and “antidote”) apply to both spores and vegetative cells.

### Sporadic toxicity affects lots of agar from multiple different vendors and across multiple decades

During a different research project in a separate laboratory, we also encountered unexpected, sudden-onset reductions in the plating efficiency of fission yeast cells, this time on Edinburg minimal media agar (EMMA). As with the process described above, we systematically tested various components of that media for potential insufficiencies or toxicities. For example we tested, one variable at a time, all combinations of three different types of water and three types of agar (**Figure 5**). Each type of water supported growth (**Figure 5**, two left columns), demonstrating that neither the water nor the nutrient components harbored any toxins; two types of agar supported growth (**Figure 5**, two left columns); and one type of agar strongly suppressed the efficiency of plating (**Figure 5**, right column). We conclude that toxicity occurs sporadically in a subset of agar lots from multiple different suppliers. Notably, we discovered the toxicity not only in agar lots from different vendors but also in lots whose purchase was widely separated in time (e.g., 2012 for data in **Figure 5** and 2023 for **Figure 4**). This time frame provides compelling evidence that those separate instances of toxicity were of independent origin.

**Figure 5.**
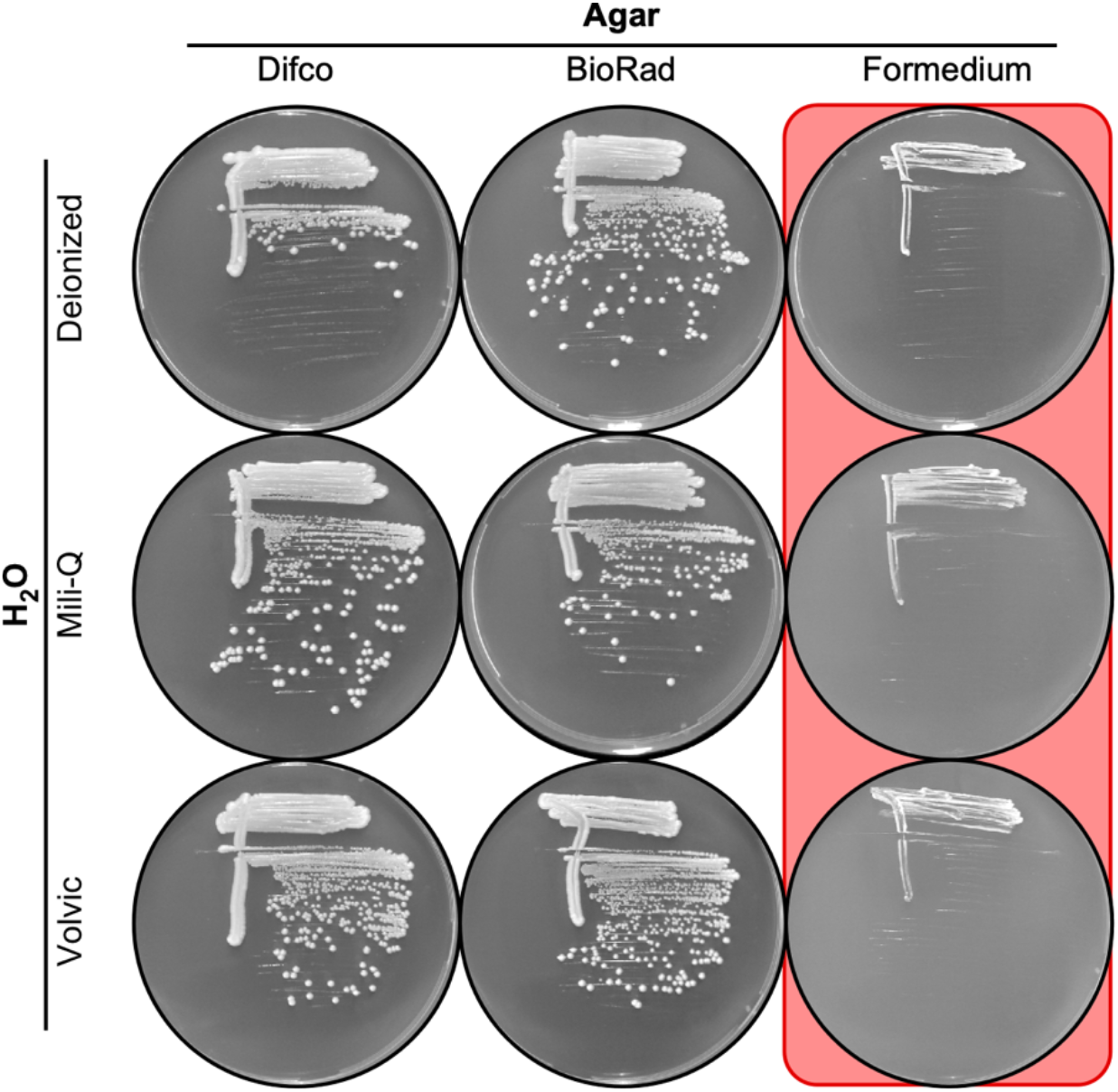
The toxicity affects agar from multiple suppliers. Nine different batches of EMMA media were prepared using the indicated lots of water and agar as the variables. Images show plating of the same strain streaked out on the different media. Note that each of the nutrient components and type of H_2_O supports growth (columns 1, 2) and that one lot of agar inhibits growth (column 3).

Our searches of scientific publication databases failed to identify papers related to adverse impacts of agar on the culture of fission yeast. Therefore, to share an advisory and to seek feedback on the potential extent of the problem, we sent a query to members of the fission yeast research community via the PombeList email server. Multiple respondents reported encountering identical, similar, or related problems with agar. Interestingly, the reported issues occurred in each of three consecutive decades and involved agar from multiple different vendors. For example, in 2006 there was apparently a batch of agar toxic to *S. pombe* that went from one wholesale source to contaminate the stocks of multiple different vendors (Nick Rhind, personal communication). Sunrise Science Products, whose agar is discussed in this article, was reportedly instrumental in identifying the toxic batch and supplying a non-toxic alternative.

In summary, our results and unpublished findings shared with us by other researchers indicate that the sporadic contamination of agar by toxic agents is relatively common and has occurred repeatedly over multiple decades of agar production.

### Rich media attenuates toxicity of the agar

The toxicity of agar can be suppressed partially by some lots of nitrogen base, with variable penetrance (**Figures 2-4**), suggesting that some lots of nitrogen base have a more abundant counteracting factor (“antidote”) than others. To see if the richness the nutrients might be responsible, we replaced the nitrogen base and ammonium sulfate (NB) with yeast extract (YE) (**Figure 6**). This change greatly suppressed—but did not counteract entirely—the toxicity of the agar, relative to the EOP for controls with nontoxic agar. For example, on toxic agar of lot 2H2428 the EOP was near zero for media whose nutrient was nitrogen base lot SLBR5876V (**Figure 6**, bottom row) but the EOP was about 35% for media whose nutrient was yeast extract lot 23C1556871 (**Figure 6**, top row). The broader implications are striking: Quality control tests that employ only the ability of fission yeast spores or cells to form colonies when streaked or spread on rich media can give false-positive assurances as to the quality of the agar; correspondingly, such tests can yield false-negative conclusions for the presence of toxicity.

**Figure 6.**
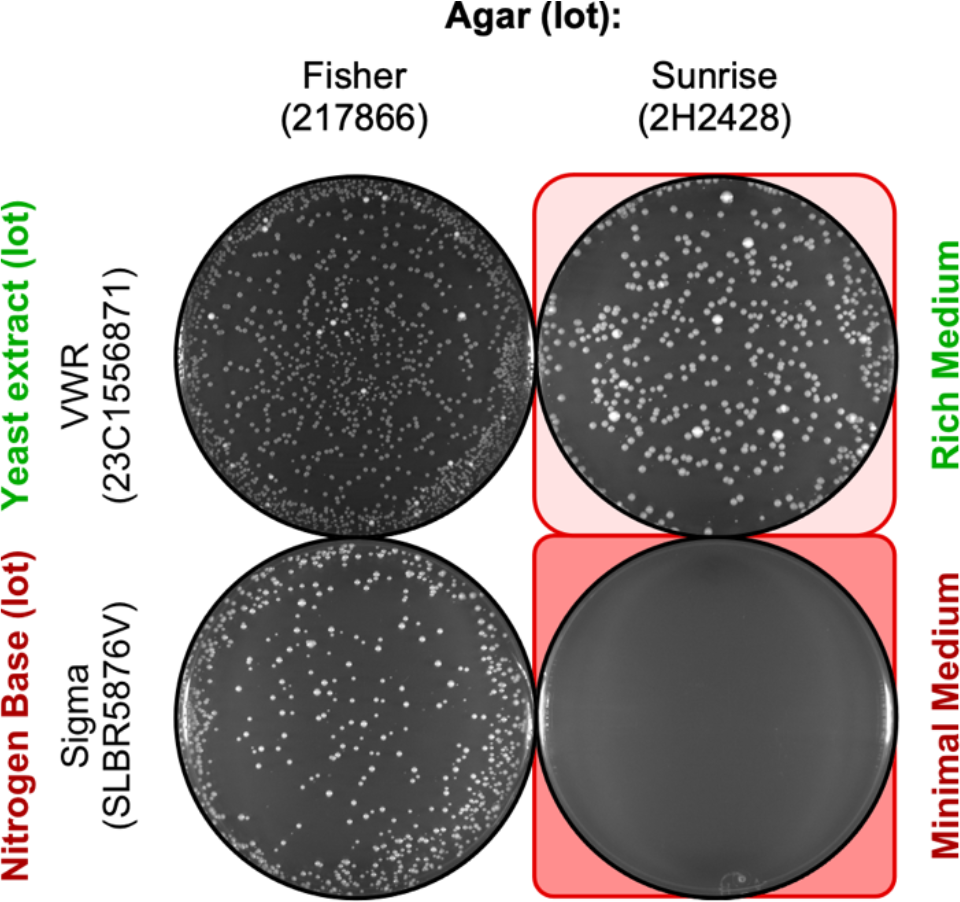
Rich media attenuates but does not abolish the agar lot-specific reduction in plating. Rich YEA (yeast extract) plates and minimal NBA plates were prepared using the indicated lots of agar. In each experiment, 2,500 spores were plated. Representative examples shows agar type-specific, reduced spore plating efficiency on each type of media.

## Discussion

### Characteristics, costs, and challenges of toxic agar

This study revealed that some lots of agar contain a yet-unidentified agent that strongly inhibits the growth of fission yeast colonies (**Figures 1-6**). The inferred toxic agent suppresses the plating efficiency of both spores and vegetative cells (e.g., **Figure 2** and **Figure 4**). The toxicity is manifest in multiple different lots of agar from the same vendor (e.g., **Figure 1** and **Figure 2**), in lots of agar from different vendors (**Figure 4** versus **Figure 5**), and in agar lots manufactured in each of three different decades (e.g., **Figure 4** and **Figure 5**; Nick Rhind, personal communication). These findings support an overarching conclusion: the sporadic, independently arising toxicity of some agar lots towards *S. pombe* is fairly common. This concern is not unique to those who culture fission yeast because specific types of agar or components therein can also affect adversely the growth of other microorganisms [e.g., (Bosmans et al., 2016; Costa & Kuramae, 2021; Rygaard et al., 2017; Tanaka et al., 2014)].

The sporadic toxicity of agar has substantial costs beyond the value of the agar itself. In our case, the reduction in EOP caused by the toxic agars interrupted our research program, which relies heavily on quantitative plating of fission yeast cells and spores. Similar concerns apply for other research and biomedical applications of agar. For example, agar toxicity might contribute to false-positive results for assays on the potency of new antibiotics or to false-negative results when testing the motility of pathogens. Similarly, while the US Food and Drug Administration and other regulatory agencies list agar as being generally recognized as safe (GRAS), it is possible that a sporadic contaminant which is potent enough to affect adversely the growth of bacterial and yeast cells might also affect mammalian cells.

Our findings revealed two main challenges to identifying and tracing toxicity: First, the problem is sporadic and not attributable to a specific vendor (e.g., **Figures 1, 4, 5**). This is not surprising, given that the production of agar is a world-wide endeavor that has historically relied on many small-scale enterprises (individuals or groups) harvesting many different species from wild sources of seaweed (Porse & Rudolph, 2017). Similarly, the manufacture of the agar is decentralized, with many producers and distributors. More fundamentally, the quality of agar is affected by high heterogeneity of numerous factors, including species, growing environments, harvesting and extraction methods, post-extraction treatments, complexity of carbohydrates, their diverse modifications, and contaminants such as fatty acids, phycobiliproteins, pigments, and secondary metabolites [see (Li & Liu, 2022) and refs therein]. And while vendors of scientific agar seek to remove factors that can inhibit microbial growth, the manufacturing processes are proprietary (i.e., opaque), the vendors do not reveal key variables such as species source, and detailed quality control measures are typically not provided to end-users. Several of these issues could be addressed satisfactorily by simple improvements to quality controls (see next section).

The second key challenge identified by this study is that the toxicity displays variable penetrance that can be modulated by the nutrient components of the media (e.g. **Figures 2-4**) and can be suppressed substantially by rich nutrients (**Figure 6**). This challenge, like the sporadic nature of toxicity (above), complicates the process of identifying the source(s) of problems with culture media: To reveal unambiguously that a problem with the culture media is caused by the lot or batch of agar, one must systematically alter—one variable at a time—both the type of agar and the type of nutrient. Moreover, if a vendor (or an end-user) “simply” wished to test whether or not an agar lot is toxic, the choice of nutrient media would be crucial. Scoring only for colony growth on rich media—which suppresses the toxicity of the agar (**Figure 6**)—can yield false-negative results for the detection of toxicity. Thus, for quality control tests to be valid, they would have to be conducted using media whose nutrient components permit (i.e., do not mask) the agar toxicity (see recommendations in next section).

### A call to improve quality control measures

Ultimate responsibility for the quality of scientific agar lies with the vendor. A fundamental, broadly applicable finding of this study is that for quality control tests to be valid, they must include precise, quantitative measurements of the EOP on all types of media typically employed for the organism of interest. For *S. pombe*, this would include measuring the EOP of both spores and vegetative cells on rich media (e.g., YEA) and minimal media (e.g., NBA and EMMA). On behalf of scientists world-wide, we call on vendors of scientific agar to conduct— and to document in writing the results of—rigorous, thorough, organism-specific tests for potential toxicity of each lot of agar as a pre-condition for its sale.

Lastly, we encourage members of research communities to be vigilant and proactive. If one encounters newly arising difficulty in plating a given organism, one should suspect—and test for—toxicity within the agar. This could be done expeditiously and inexpensively by comparing growth within liquid media to that on solid media. Confirmed or suspected problems with agar should be brought as soon as possible to the attention of the vendor and members of the relevant research community (e.g., via the PombeList email server). These actions will help colleagues to avoid wasting their precious time and resources and will help the vendors to identify and correct potential defects in their products.

## Acknowledgements

We thank members of the PombeList community for feedback about toxic agar; Seth Dixon and Emory Malone for comments on the manuscript. This study was supported by a research project grant from the National Institute of General Medical Sciences at the National Institutes of Health to WPW (grant number **R01 GM145834**); and by funds from the CNRS (ATIP-Avenir) and the Agence Nationale de la Recherche to DH (award number **ANR-15-CE12-0009-01**).

## Authors’ contributions

All authors contributed to experimental design. Reine Protacio, Dominique Helmlinger and Mari Davidson conducted experiments and analyzed data. Wayne Wahls constructed figures and wrote the manuscript. All authors edited and approved the final manuscript.

## Data availability statement

All data necessary to support the conclusions of this study are contained within the article. Yeast strain reagents are available from the authors upon request.

